# Melanoma plasticity is controlled by a TRIM28-JUNB mediated switch

**DOI:** 10.1101/777771

**Authors:** William A Nyberg, Tianlin He, Maria Sjöstrand, Diego A Velasquez-Pulgarin, Lucia Pellé, Ruxandra Covacu, Alexander Espinosa

**Affiliations:** Center for Molecular Medicine, Department of Medicine, Karolinska Institutet, Karolinska University Hospital, Stockholm, Sweden; Center for Cell Engineering and Department of Medicine, Memorial Sloan-Kettering Cancer Center, New York, NY, USA; Department of Biomedical Engineering, University of Memphis, Memphis, TN, USA; Center for Molecular Medicine, Department of Clinical Neuroscience, Karolinska Institutet, Karolinska University Hospital, Stockholm, Sweden; Mosaiques Diagnostics GmbH, Rotenburger Str. 20, 30659 Hanover, Germany

## Abstract

The introduction of immune checkpoint blockade has revolutionized the treatment of metastatic melanoma^1^. However, 40-60% of patients with metastatic melanoma do not respond to immune checkpoint blockade, and a significant fraction of patients acquire resistance to treatment^2,3^. This resilience and aggressiveness of melanoma tumors is partly due to their ability to switch between invasive and proliferative states^4,5^. The transition between phenotypic states indicates that phenotype switching occurs through reversible epigenetic mechanisms rather than by acquisition of mutations^6,7^. Identifying the epigenetic mechanisms that underlie phenotype switching of melanoma cells could potentially lead to new therapeutic strategies. Here we report that the bromodomain protein TRIM28 (KAP1/TIF1β) regulates a JUNB dependent phenotypic switch in melanoma cells. Knockdown of TRIM28 in melanoma cells reduced the expression of pro-invasive YAP1 signature genes, and led to reduced invasiveness and lung colonization. In contrast, TRIM28 knockdown increased the expression of KRAS signature genes and promoted tumor growth. TRIM28 interacted with the transcriptional elongation factors CDK9 and HEXIM1, and negatively regulated the transcriptional elongation of JUNB by RNA polymerase II. Rescue experiments demonstrated that the effects of TRIM28 knockdown were directly mediated by JUNB. Mechanistically, JUNB played a pivotal role in phenotype switching by inhibiting the invasiveness of melanoma cells and increasing the growth of melanoma tumors. Our results contribute to the understanding of melanoma plasticity, and suggest that cancer drugs inhibiting the transcriptional elongation of RNA polymerase II should be carefully evaluated in melanoma to exclude the risk for increased metastasis.

---

To identify bromodomain genes associated with aggressive melanoma, we analyzed RNA-seq and whole exome sequencing data from 367 metastatic tumors together with analysis of corresponding survival data (TCGA-SKCM). Using unsupervised principal component analysis, and partitioning around medoids clustering, we identified two clusters of patients (Fig. 1a). Cluster C2 was characterized by high expression of the bromodomain genes *TRIM28, SMARCA4, BRD4*, and *BRPF1* (Fig. 1b), and survival analysis revealed that patients in this cluster also had significantly shorter overall survival (Fig. 1c). *TRIM28* was the bromodomain gene most strongly associated with poor overall survival (Fig. 1c and Extended Data Fig. 1a, b). Analysis of whole exome sequencing data did not reveal any differences in previously described melanoma mutations between cluster C1 and C2^10^ (Fig. 1d). Also, no difference in bromodomain gene mutations was found between cluster C1 and C2 (Extended Data Fig 1c). In all, we identified a subset of melanoma patients that is characterized by poor survival and a distinct bromodomain gene signature with high expression of *TRIM28*.

**Figure 1.**
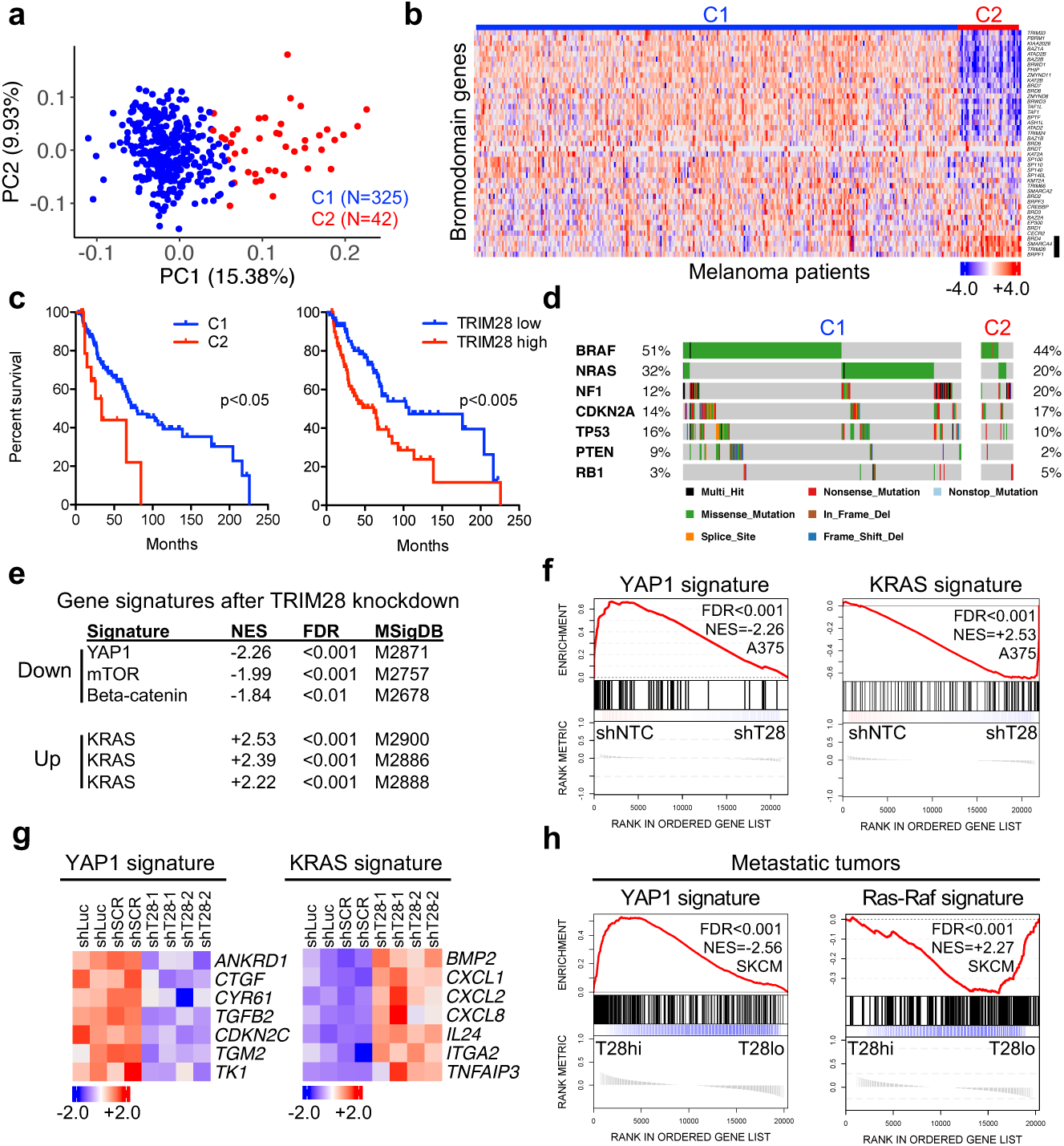
TRIM28 controls a switch between YAP1 and KRAS transcriptional programs. **a**, Principal component analysis of patients with cutaneous malignant melanoma based on global gene expression (RNA-seq) in metastases (n=367). Partitioning around medoids clustering algorithm was used for clustering (k=2). **b**, Heat-map displaying the expression of bromodomain genes in metastases from patients with cutaneous malignant melanoma (n=367). *BRD4, SMARCA4, TRIM28* and *BRPF1* are highlighted by a black bar. Gene expression is represented by z-score. **c**, Kaplan-Meier analysis of overall survival of patients with stage III melanoma in cluster C1 and C2, and of patients with stage III melanoma with high or low TRIM28 expression. The log-rank test was used for statistical testing of survival data. **d**, Frequencies of mutations in common oncogenes and tumor suppressor genes in the C1 and C2 tumor clusters. **e**, The most up- or downregulated oncogenic gene signatures after GSEA of A375 cells transduced with TRIM28 specific shRNA (shT28-1 or shT28-2) or non-targeting control (shNTC) shRNA (shLuc or shSCR). **f**, GSEA plots of YAP1 signature genes (MSigDB M2871) and KRAS signature genes (MSigDB M2900) in A375 cells transduced with two shT28 (shT28-1 or shT28-2) or two shNTC (shLuc or shSCR). **g**, Expression of canonical YAP1 and KRAS genes in A375 cells transduced with two shT28 (shT28-1 or shT28-2) or two shNTC (shLuc or shSCR). Gene expression is represented by z-score. **h**, GSEA plots of YAP1 signature genes^22^ and the Ras-Raf gene signature (MSigDB M2728) in metastases from patients with cutaneous malignant melanoma. Data used for analysis in (**a-d, h**) were downloaded from The Cancer Genome Atlas. Statistical testing of GSEA (**e, f, h**) was performed as described in Methods. GSEA, gene set enrichment analysis; FDR, false discovery rate; NES, normalized enrichment score, MSigDB, Molecular Signature Database.

TRIM28 is a transcriptional regulator that is necessary for the repression of transposable elements, the maintenance of epigenetic stability and for the control of transcriptional elongation by RNA polymerase II^11-13^. Due to its roles in regulating gene expression, we hypothesized that TRIM28 controls oncogenic transcriptional programs in melanoma cells. To test this, we transduced A375 cells with two TRIM28 specific shRNAs (shT28-1 or shT28-2), and two non-targeting control shRNAs (shNTC). Knockdown efficiency was verified using immunoblotting and quantitative RT-PCR (Extended Data Fig. 2a, b). Using gene set enrichment analysis (GSEA) we identified YAP1 regulated genes as the most repressed oncogenic signature after TRIM28 knockdown, and genes downstream of Ras signaling (KRAS signature) as the most increased oncogenic signature (Fig. 1e-g). These results were verified in independently generated TRIM28-knockdown A375 cells, and in TRIM28-knockout A375 cells generated by CRISPR/Cas9 (Extended Data Fig. 2c-f). We then asked if TRIM28 was associated with the expression of YAP1 and KRAS signature genes in metastases from melanoma patients. Indeed, we found that low TRIM28 expression in metastatic tumors was associated with low expression of YAP1 signature genes and with high expression of KRAS signature genes (Fig. 1h). In addition, TRIM28 knockdown cells and metastatic tumors with low TRIM28 expression had reduced expression of metastatic gene signatures (Extended Data Fig. 4a-c). To determine if TRIM28 regulated Ras signaling, we measured the phosphorylation of 24 signaling mediators (MAPK/ERK and PI3K/AKT/mTOR pathways) in shSCR and shT28-1 transduced A375 cells. We found no evidence of altered signaling activity after TRIM28 knockdown in either A375 and A2058 cells (Extended Data Fig. 3a-c). These results showed that TRIM28 controls a switch between the YAP1 (invasive) and KRAS (proliferative) transcriptional programs in melanoma.

The transcription factor YAP1 promotes migration and invasiveness of melanoma cells^14,15^, while KRAS signature genes promote proliferation and tumor growth^16^. We therefore hypothesized that TRIM28 is necessary for the metastatic potential of melanoma cells and for controlling the growth of melanoma tumors. To test the role of TRIM28 in melanoma invasiveness, we first performed intravenous injections of A375-MA2 cells stably expressing shT28-1, shT28-2 or scrambled shRNA (shSCR). Eight weeks after injection, all mice injected with shSCR transduced A375-MA2 cells had lung tumors (Fig. 2a, b). In contrast, only one mouse injected with shT28-1 or shT28-2 transduced A375-MA2 cells had lung tumors (Fig. 2a, b). We then performed Matrigel invasion assays, and found that knockdown of TRIM28 or YAP1 resulted in reduced invasiveness of A375 cells (Fig. 2c, d). TRIM28 has previously been reported to promote growth of breast and prostate cancer^8,9^. However, the increased levels of the mitogenic and angiogenic factors CXCL1, CXCL2 and CXCL8^17^ after TRIM28 knockdown suggested that TRIM28 instead suppresses the growth of melanoma tumors. To test this, we performed subcutaneous injections of A375 cells stably transduced with shT28-1 or shSCR lentivirus. Indeed, in contrast to its role in breast and prostate cancer cells^8,9^, knockdown of TRIM28 in melanoma cells led to more rapid tumor growth (Fig. 2e, f). We verified this result using an alternative TRIM28-specific shRNA construct, and also with Trim28 knockdown in mouse B16.F10 melanoma cells (Extended Data Fig. 5a-d). The metastatic capacity of melanoma cells is linked to drug resistance, and this combination is the main cause of mortality in malignant melanoma^18^. We therefore asked if TRIM28 knockdown affected the drug sensitivity of A375 and A2058 cells, and found that both TRIM28 and YAP1 knockdown increased the sensitivity to the alkylating agents Dacarbazine and Temozolomide (Extended Data Fig. 6a, b). Taken together, these results showed that TRIM28 controls a phenotypic switch between invasiveness and tumor growth of melanoma. Specifically, TRIM28 was necessary for the invasiveness of melanoma cells while simultaneously suppressing tumor growth.

**Figure 2.**
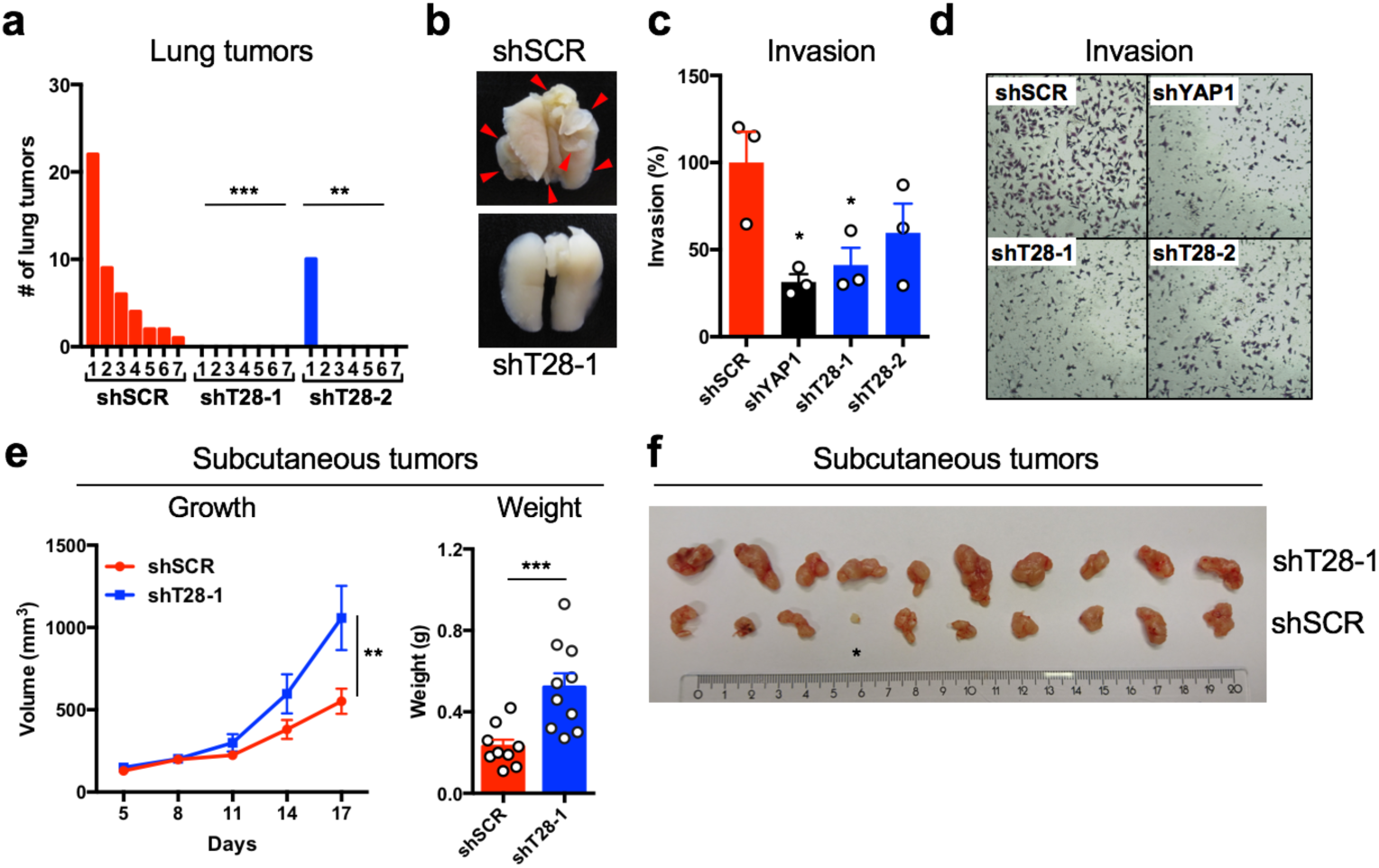
TRIM28 controls a phenotypic switch between invasiveness and tumor growth in melanoma. **a**, Female nu/nu BALB/c mice were injected intravenously with 1.5×10^5^ A375-MA2 cells stably transduced with scrambled control shRNA (shSCR) or TRIM28 specific shRNAs (shT28-1 or shT28-2) (n=7 mice per shRNA). Counting of lung tumors was performed in a blinded manner. One representative experiment of two is shown. One-way ANOVA and Tukey’s post hoc test was used for statistical testing of tumor numbers. **b**, Representative images of lungs from (**a**). **c**, Quantification of Matrigel invasion experiments with A375 cells transduced with shSCR, shT28-1, shT28-2, or a YAP1 specific shRNA (shYAP1). Results are expressed as mean ±s.e.m. from three independent experiments (n=3). One-way ANOVA and Tukey’s post hoc test was used for statistical testing. **d**, Representative images from one of three Matrigel invasion assays from (**c**). **e**, Tumor growth after subcutaneous injection of 2.5×10^6^ A375 cells transduced with shSCR or shT28-1 lentivirus (n=10 mice per group). The tumor weight was analyzed 17 days after subcutaneous injection. Results are expressed as mean ±s.e.m. One experiment of two is shown. Repeated-measures ANOVA was used for the statistical test of tumor growth, and the two-sided Mann-Whitney U-test was used for the statistical test of tumor weight. **f**, Picture of tumors 17 days after subcutaneous injection of cells transduced with shSCR or shT28-1. The tumor marked * was excluded from data analysis due to lack of engraftment.

To elucidate how TRIM28 regulated the phenotypic switch between invasiveness and proliferation, we mapped the TRIM28 interactome in A375 cells using biotin-based proximity proteomics (Fig. 3a). We identified several chromatin-associated proteins involved in epigenetic regulation of gene expression and RNA polymerase II (RNAPII) transcriptional elongation, including SETDB1, CBX5, CDK9 and HEXIM1 (Fig 3b and Extended Data Fig. 7a-c). As expected, we also identified >50 KRAB-ZFN proteins as interactors of TRIM28^19,20^, validating our proximity proteomics approach. As TRIM28 knockdown led to increased expression of several immediate early genes, including *JUNB, FOSL1* and *MYC*, this indicated that TRIM28 negatively regulated transcriptional elongation by RNAPII (Fig 3c). We analyzed ChIP-seq data and found significant genome-wide overlap between TRIM28, CDK9 and HEXIM1, with a common peak around 200 bp downstream of the transcription start site (TSS) (Extended Data Fig. 7d). We also identified overlapping ChIP-seq signals of TRIM28, CDK9, HEXIM1, and RNAPII in the immediate early genes *FOSL1* and *JUNB* (Fig. 3d).

**Figure 3.**
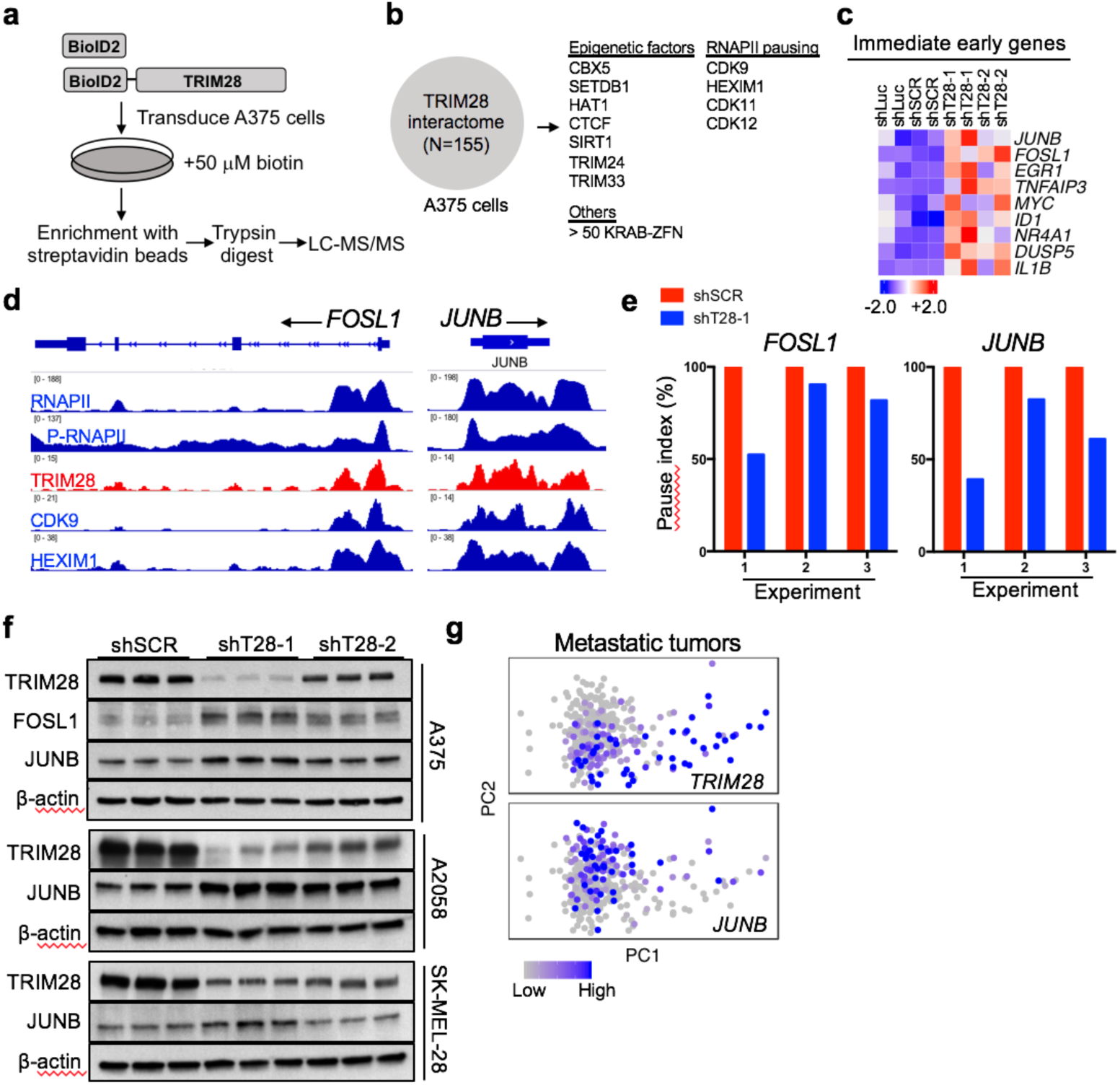
TRIM28 negatively regulates the transcriptional elongation of *FOSL1* and *JUNB*. **a**, A375 cells were transduced with pBABE-BioID2-TRIM28 or pBABE-BioID2 retrovirus and selected with 1μg/ml puromycin. Transduced cells were cultured in complete DMEM medium containing 50 μM biotin for 24 hours, followed by enrichment of biotinylated proteins using streptavidin beads, and identification of interactors with LC-MS/MS. **b**, Identified TRIM28 interactome in A375 cells. **c**, Expression of genes regulated by RNAPII transcriptional elongation in A375 cells transduced with shNTC or shT28 lentivirus. Gene expression is represented by z-score. **d**, ChIP-seq analysis of co-occupancy of TRIM28, CDK9, HEXIM1 and RNAPII at *FOSL1* and *JUNB* in HCT-116 cells. **e**, RNAPII pausing index for *FOSL1* and *JUNB* genes in A375 cells transduced with shSCR or shT28-1 lentivirus. The pausing index was calculated in serum starved cells. Results from three independent experiments (n=3) are shown. **f**, Protein levels of TRIM28, FOSL1 and JUNB in A375 cells transduced with shSCR, shT28-1 or shT28-2 lentivirus. One representative experiment of two is shown. **g**, Expression levels of *TRIM28* and *JUNB* are indicated for metastatic tumors (TCGA-SKCM), and displayed in a principal component analysis plot (n=367). LC-MS/MS, liquid chromatography-mass spectrometry; RNAPII, RNA polymerase II; P-RNAPII, phospho-RNA polymerase II.

To verify that TRIM28 negatively regulates transcriptional elongation in melanoma cells, we determined the extent of RNAPII pausing of *JUNB* and *FOSL1* in A375 cells stably expressing shT28-1 or shSCR. After ChIP with an anti-RNAPII antibody, quantitative PCR was performed to quantify the precipitated genomic regions of *JUNB* and *FOSL1*. A pausing index was calculated for each gene by dividing the RNAPII occupancy at the TSS with the RNAPII occupancy in the gene body (+1 kb). We observed a decreased pause index for both *JUNB* and *FOSL1* after TRIM28 knockdown, indicating that TRIM28 prevents RNAPII release and thus negatively regulates transcriptional elongation (Fig. 3f). Verifying the reduced RNAPII pausing of *JUNB* and *FOSL1*, we found increased protein levels of FOSL1 and JUNB after TRIM28 knockdown in A375, A2058 and SK-MEL-28 cells (Fig. 3g). In addition, low *TRIM28* expression in metastases from melanoma patients was associated with high *JUNB* expression (Fig. 3h). Taken together, these results show that TRIM28 controls the expression of *FOSL1* and *JUNB* by negatively regulating the transcriptional elongation of RNAPII.

JUN/FOS transcription factors play dual roles in cancer cells by controlling the expression of both YAP1 target genes and genes downstream of Ras signaling^21^. We therefore asked if FOSL1 and JUNB controls the expression of YAP1 and KRAS signature genes in melanoma cells. To test this, we first overexpressed FOSL1 or JUNB in A375 and A2058 cells. We found that JUNB was sufficient to suppress YAP1 signature genes (*ANKRD1, TGFB2, CDKN2C*) and to activate KRAS signature genes (*CXCL2, CXCL8*) (Fig. 4a, b). In contrast, FOSL1 overexpression increased the expression of YAP1 signature genes, but was insufficient to activate the expression of *CXCL2* and *CXCL8* (Extended Data Fig. 8a, b). We then used CRISPR interference (CRISPRi) to validate the role of JUNB in suppressing YAP1 signature genes. Indeed, reduced JUNB levels led to increased expression of YAP1 target genes, revealing a necessary role for JUNB in suppressing the YAP1 transcriptional program in melanoma cells (Fig. 4c). To verify that the transcriptional changes observed in TRIM28 knockdown cells were mediated by JUNB, we performed a rescue experiment by transfecting TRIM28 knockdown cells with non-targeting siRNA (siNTC) or JUNB-targeting siRNA (siJUNB). Indeed, transfection with siJUNB restored the expression of both YAP1 and KRAS signature genes (Extended Data Fig. 9a-d). In all, these results demonstrated that JUNB was necessary and sufficient to mediate the changes in YAP1 and KRAS gene signatures observed after TRIM28 knockdown.

**Figure 4.**
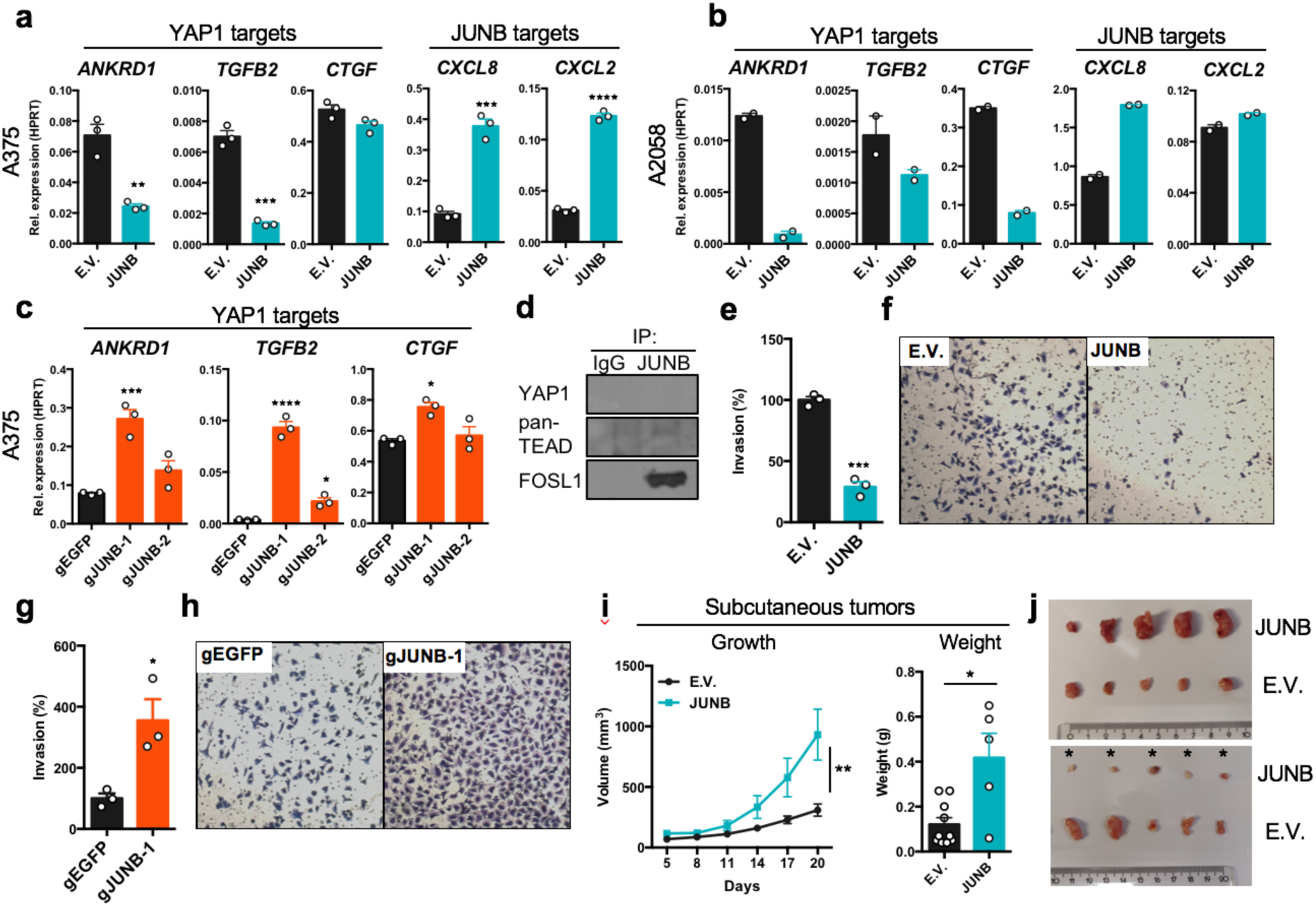
JUNB mediates a switch between melanoma invasiveness and growth. **a**, Expression of YAP1 or JUNB target genes in A375 cells after transduction with pBABE-JUNB or pBABE-E.V. (empty vector). Results are expressed as mean ±s.e.m. from three independent experiments (n=3). Two-sided unpaired t-tests were used for statistical testing. **b**, Expression of YAP1 or JUNB target genes in A2058 cells after transduction with pBABE-JUNB or pBABE-E.V. (empty vector). Results are expressed as mean ±s.e.m. from two independent experiments (n=2). **c**, Expression of YAP1 target genes in A375 cells after transduction with dCas9-KRAB lentiviruses expressing gRNA targeting *JUNB* (gJUNB-1 or gJUNB-2) or a control gRNA (gEGFP). Results are expressed as mean ±s.e.m. from three independent experiments (n=3). One-way ANOVA and Dunnett’s multiple comparison test were used for statistical testing. **d**, Immunoprecipitation of endogenous JUNB in A375 cells followed by immunoblotting against YAP1, pan-TEAD and FOSL1. **e**, Quantification of Matrigel invasion with A375 cells transduced with pBABE-JUNB or pBABE-E.V. (empty vector). Results are expressed as mean ±s.e.m. from three independent experiments (n=3). Two-sided unpaired t-tests were used for statistical test between the groups. **f**, Representative images from one of three Matrigel invasion assays. **g**, Quantification of Matrigel invasion experiments with A375 cells transduced with dCas9-KRAB lentiviruses expressing gRNA targeting *JUNB* (gJUNB-1 or gJUNB-2) or a control gRNA (gEGFP). Results are expressed as mean ±s.e.m. from three independent experiments (n=3). Two-sided unpaired t-tests were used for statistical test between the groups. **h**, Representative images from one of three Matrigel invasion assays. **i**, Tumor growth after subcutaneous injection of 2.0×10^6^ A375 cells transduced with JUNB or E.V. lentivirus (n=10 mice per group). The tumor weight was analyzed 20 days after subcutaneous injection. Results are expressed as mean ±s.e.m. Repeated-measures ANOVA was used for the statistical test of tumor growth, and the two-sided Mann-Whitney U-test was used for the statistical test of tumor weight. **j**, Picture of tumors 20 days after subcutaneous injection of cells transduced with JUNB or E.V. lentivirus. Tumors marked * were excluded from data analysis due to lack of engraftment.

In contrast to other JUN/FOS transcription factors^21-23^, JUNB has not been reported to bind YAP1/TEADs or to control the YAP1 transcriptional program. We therefore asked if JUNB inhibited the expression of YAP1 signature genes by binding to YAP1 or TEADs in melanoma cells. Co-immunoprecipitation against endogenous JUNB did not reveal any interaction to YAP1 or TEADs, instead we detected an interaction between JUNB and FOSL1 (Fig. 4d). The genome-wide overlap between JUNB and TEAD4 peaks therefore suggests that JUNB does not suppress YAP1 signature genes by binding to YAP1/TEADs, but instead by binding to the YAP1/TEAD co-activator FOSL1 (Fig 4e and Extended Data Fig. 9e).

The role of JUNB in suppressing YAP1 transcriptional program in melanoma cells suggested that JUNB negatively regulates melanoma invasiveness. To test this, we performed Matrigel invasion assays and found that overexpression of JUNB significantly reduced the invasiveness of A375 cells (Fig. 4f, g). Conversely, CRISPRi against JUNB led to a flattened mesenchymal-like cell morphology and increased melanoma invasiveness (Fig. 4h, i, and Extended Data Fig 10a). Targeting JUNB with CRISPRi also led to increased expression of *MITF, TYR* and the invasive marker *CDH2* (N-cadherin) (Extended Data Fig 10b-g). Consistent with its role in reducing invasiveness and promoting growth, overexpression of JUNB led to reduced tumor engraftment and increased growth of established tumors (Fig. 4j, k). Taken together, our data show that JUNB suppresses invasiveness and promotes tumor growth by controlling the expression of YAP1 and KRAS signature genes. JUNB thus plays a pivotal role in melanoma plasticity.

The plasticity of melanoma cells underlies their ability to metastasize and to develop drug addiction^24^. Here we have uncovered a new TRIM28-JUNB mediated phenotypic switch in melanoma cells. TRIM28 controlled the transcriptional elongation of JUNB, that then regulated transcriptional programs for invasiveness and tumor growth. Since the transcriptional elongation by RNA polymerase II has emerged as a promising target for cancer drugs^25,26^, we propose that such drugs should be carefully evaluated in melanoma to exclude the risk for increased metastasis.

## METHODS

### Analysis of patient data

We downloaded RNA-seq data, mutation data, and clinical data, from patients with metastatic melanoma (TCGA-SKCM, https://portal.gdc.cancer.gov/). To analyze RNA-seq data from metastatic tumors of patients with survival data (N=367), we first filtered gene expression (TPM) using the *genefilter* package in R using pOverA(p=0.75, A=100). We then used the *ggplot2* and *ggfortify* packages in R to perform principal component analysis. The *factoextra* package in R was used to determine the optimal numbers of clusters, and partitioning around medoids (PAM) clustering (k=2) was performed using the *cluster* package in R. Cox regression analysis and Kaplan-Meier survival analysis was performed using the *survival* package in R and in Prism 6 (GraphPad Software). For mutational analysis, we downloaded MAF files from whole exome sequencing of metastatic melanoma tumors (TCGA-SKCM), and used the *maftools* package in R for mutational analysis and visualization.

### Human Transcriptome Array 2.0

A375 cells were transduced with lentiviruses expressing non-targeting control shRNAs (shSCR or shLuc), or TRIM28 specific shRNAs (shT28-1 or shT28-2), followed by selection with puromycin (1 μg/ml). Cells were harvested in cold PBS and lysed in RLT buffer (Qiagen). RNA quality and integrity were verified with the Bioanalyzer system (Agilent). The GeneChip Human Transcriptome Array (HTA) 2.0 (Affymetrix) was used for determination of global gene expression. RNA quality control and HTA 2.0 microarray analysis was performed at the Bioinformatics and Expression Analysis core facility at Karolinska Institutet. CEL files from the HTA 2.0 microarrays were preprocessed and normalized with robust multi-array average (RMA) using the R package *oligo*. Data were log_2_ transformed and normalized to z-scores, and heatmaps were generated using the R package *ComplexHeatmap*. Significance Analysis of Microarrays (SAM) analysis of gene expression was performed using the R packages *siggenes* and *genefilter.* Volcano plots were generated using the R packages *ggplot2, ggrepel* and *gghighlight*.

### Gene set enrichment analysis

Expression data files (.gct), phenotype labels (.cls), and gene set files (.gmx) were uploaded to Genomspace (http://www.genomespace.org/) and analyzed using the GSEA tool in GenePattern^27^. RNA-seq data were analyzed using unfiltered natural scale values. Microarray data were analyzed using log_2_ transformed values. Normalized enrichment scores and false discovery rates were calculated as described^27^. Batch GSEA was performed for oncogenic signatures (Molecular Signatures Database v6.2, 189 gene sets), while metastatic gene signatures were analyzed with selected gene sets. Enrichment plots were generated using the *replotGSEA* function in R (https://github.com/PeeperLab/Rtoolbox).

### ChIP-seq analysis

BigWig files for selected transcription factors were downloaded from the Gene Expression Omnibus repository (https://www.ncbi.nlm.nih.gov/geo/) and ChIP-seq tracks were visualized using the Integrated Genomics Viewer (IGV)^28^. The following data sets were used for ChIP-seq analysis: GSE72622 (CDK9, HEXIM1, TRIM28, RNAPII, and Phospho-RNAPII), GSE32465 (TEAD4) and GSE92807 (JUNB). Profile plots around TSS were generated in Galaxy (http://usegalaxy.org) using the computeMatrix tool^29^. We used the R package *ChIPpeakAnno* for determining genome wide overlap between transcription factors.

### Cell culture

Cell lines used in this study were: HEK-293T (Espinosa laboratory stock), A375 (American Type Culture Collection), A375-MA2 (American Type Culture Collection), SK-MEL-28 (American Type Culture Collection), A2508 (American Type Culture Collection) and B16-F10 (American Type Culture Collection). All cell lines were routinely tested for mycoplasma contamination. Cells were cultured in high-glucose DMEM (Sigma Aldrich) supplemented with fetal calf serum (10%), streptomycin (0.1 mg/ml), penicillin (100 U/ml), Sodium pyruvate (1 mM) (Sigma Aldrich), HEPES (10 mM) (Sigma Aldrich) and L-glutamine (2 mM) (Sigma Aldrich). All viruses were packaged for 48 hours following transfection of HEK-293T cells using X-tremeGENE 9 DNA Transfection Reagent (Roche).

### Animal studies

Eight-week-old female nude mice (BALB/cAnNRj-Foxn1^nu/nu^, Janvier Labs) were injected subcutaneously with 2.0×10^6^ (JUNB experiment) or 2.5×10^6^ (shT28 experiments) A375 cells in 100 μl of Matrigel (#354263, Corning) diluted to 50% in PBS. Tumor size was measured every third day using a digital caliper, and tumor size was calculated using the formula V=(LxWxW)/2. After termination of the xenograft experiments, the tumors were weighed and biopsied for RNA extraction. For lung colonization experiments, six- to eight-week-old female nude mice (BALB/cAnNRj-Foxn1^nu/nu^, Janvier Labs) were injected intravenously (tail vein) with 1.5×10^5^ A375-MA2 cells in 100 μl PBS. The animals were sacrificed after 8 weeks and lungs were collected and fixed for 48 hours in 4% paraformaldehyde before being analyzed for lung tumors in a dissecting microscope. Lung tumor counts were performed in a blinded manner. Eight-week-old female C57BL6/J were injected subcutaneously with 1×10^5^ B16.F10 cells in 200 μl of Matrigel (#354263, Corning) diluted to 50% in PBS. After termination of the experiments the tumors were weighed and biopsied for RNA extraction. Mice were housed in a specific pathogen-free animal facility at Center for Molecular Medicine, Karolinska Institutet. The study was approved by the Ethical Review Committee North, Stockholm County, and animals were handled in compliance with the guidelines at Karolinska Institutet.

### Matrigel invasion assay

2.5×10^5^ A375 cells were seeded in 0.1% FBS DMEM in 24-well Matrigel GFR Invasion Chambers (#734-1049, Corning). The inlets were put in wells containing 10% FBS DMEM and incubated for 24 hours. The cells were fixed in buffered 4% paraformaldehyde, permeabilized with 100% methanol, and stained using Crystal Violet. For each inlet, five images were obtained randomly using a Nikon TMS-F microscope with a DeltaPix camera module, and cells were counted and averaged for each inlet using ImageJ.

### TRIM28 interactome analysis

A375 cells were transduced with pBABE-BioID2-TRIM28 or pBABE-BioID2 followed by selection with 1 μg/ml puromycin. Transduced cells were then cultured in the presence of 50 μM Biotin (Sigma Aldrich) for 20 hours prior to lysis and then bound to Dynabeads MyOne Streptavidin C1 (Thermo Fisher Scientific) magnetic beads overnight. We then performed streptavidin pull-down of biotinylated proteins following the procedure described by Roux et al.^30^. After enrichment of biotinylated proteins, they were on-bead digested using trypsin. Peptides were dissolved in 25 µl of 2% acetonitrile, 0.1% formic acid. 10% of the sample was analyzed by nanoLC-MS/MS using an Easy-1000 nLC chromatographic system (Thermo Fisher Scientific) coupled to a Q Exactive Plus mass spectrometer (Thermo Fisher Scientific). The peptides were separated using a heated (55°C) 50 cm C-18 Easy-column (Thermo Fisher Scientific), and the separation was performed using an acetonitrile/water gradient (buffer A: 2% acetonitrile, 0.1% formic acid; buffer B: 98% acetonitrile, 0.1% formic acid) of 4-26% B over 120 minutes, followed by a 26-95% B over 5 minutes and 95% B for 8 minutes. The flow rate was 300 nl/min. The instrument was operated in a data-dependent mode selecting the 16 most intense precursors in the survey mass scans at 140,000 and followed by MS/MS data acquisition at 70,000 mass resolution using Higher-energy collisional dissociation (HCD) fragmentation. Protein identification and quantification was performed at the Proteomics Biomedicum core facility, Karolinska Institutet. To discriminate genuine interactors from contaminating proteins and non-specifically bound proteins, we filtered identified proteins using the CRAPome contamination repository (http://crapome.org/). Network analysis of identified proteins was then performed using Ingenuity Pathway Analysis software (Qiagen).

### Plasmid constructs

To express short-hairpin RNA (shRNA) we used the following lentiviral plasmids: pLKO.1-TRIM28-1 (TRCN0000018001), pLKO.1-TRIM28-2 (TRCN0000018002), pLKO.1-TRIM28-3 (TRCN0000017998), pLKO.1-Trim28 (TRCN0000071366), and pLKO.1-YAP1 (TRCN0000107266). As non-targeting controls, we used pLKO.1 encoding scrambled or luciferase specific shRNA. For CRISPR/Cas9, we inserted a TRIM28 specific gRNA sequence (GGCCCCCGGCGGCGTGTGAA) into pLentiCRISPRv2 (#52961, Addgene). For CRISPRi we inserted gRNA sequences into pLV-hU6-sgRNA-hUbC-dCas9-KRAB-T2a-Puro (#71236, Addgene). The following gRNA sequences were used for CRISPRi: gJUNB-1 TAGCGCGGTATAAAGGCGTG, gJUNB-2 CCAATCGGAGCGCACTTCCG and gEGFP GACCAGGATGGGCACCACCC. To generate FOSL1 expressing retrovirus, the full coding-sequence of FOSL1 was cloned by PCR and inserted into pFLAG-CMV-6c, followed by transfer of the FLAG-FOSL1 fragment into the pBABE-puro backbone. To generate JUNB expressing retrovirus, FLAG-JUNB was excised from pCS2-FLAG-JUNB (#29687, Addgene) and inserted into the pBABE-puro backbone. To identify the TRIM28 interactome in melanoma cells, the coding sequence for TRIM28 was inserted into the pBABE-BioID2 plasmid (#80900, Addgene). To diminish the risk for steric hindrance, and to increase the radius of the interactome, a flexible linker (GGGGS) was inserted between the BioID2 tag and TRIM28. All lentiviruses were packaged using pMD2.G (#12259, Addgene) and psPAX2 (#12260, Addgene). Retroviruses were packaged using pMD2.G (#12259, Addgene) and pUMVC (#8449, Addgene).

### Quantitative PCR

All RNA extractions were performed using TRIzol (Invitrogen). cDNA conversions were performed using the High Capacity cDNA Reverse Transcription Kit (Applied Biosystems) or the iScript cDNA Synthesis Kit (Bio-Rad). Gene expression was determined using TaqMan gene expression assays (Thermo Fisher Scientific): *TRIM28* (Hs00232212_m1), *Trim28* (Mm00495594_m1), *ANKRD1* (Hs00173317_m1), *CYR61* (Hs00998500_g1), *CTGF* (Hs00170014_m1), *FOSL1* (Hs00606343_g1), *JUNB* (Hs00357891_s1), *CXCL1* (Hs00236937_m1), *CXCL2* (Hs00601975_m1), *CXCL8* (Hs00174103_m1), *TGFB2* (Hs00234244_m1), *YAP1* (Hs00902712_g1), *CDH2 (*Hs00983056_m1), *MITF* (Hs01117294_m1), *TYR* (Hs00165976_m1), *HPRT1* (Hs01003267_m1), *Hprt1* (Mm01545399_m1), *ACTB* (Hs01060665_g1) and *Gapdh* (Mm00484668_m1). Precipitated genomic sequences after ChIP in RNAPII pausing experiments were detected using SYBR based quantitative PCR using the following primers: FOSL1-TSS (F: GTGGTTCAGCCCGAGAACTT, R: AGTCTCGGAACATGCCCG). FOSL1-1kb (F: CTGTCGAGGGGCTGCG, R: CCGTGACTCGGCGGAAC). JUNB-TSS (F: GGCTGGGACCTTGAGAGC, R: GTGCGCAAAAGCCCTGTC). JUNB-1kb (F: CATCAAAGTGGAGCGCAAG, R: TTGAGCGTCTTCACCTTGTC).

### RNAPII pausing assay

A375 cells were starved for 48 hours using serum-free DMEM medium. Cells were then collected without, or 30 minutes after, addition of 10% serum. 3×10^6^ cells were collected and lysed per sample. ChIP was performed using the MAGnify Chromatin Immunoprecipitation System (Invitrogen) following the manufacturer’s protocol. Chromatin shearing was performed using a Bioruptor UCD-200 sonicator running 40 cycles of 30 seconds on and 30 seconds off at high intensity (at 4°C). For each immunoprecipitation, 3 μg anti-RNA polymerase II (ab817 Abcam) antibody was added to 10 μl chromatin. Input controls were included without addition of antibodies. Quantitative PCR was performed on purified genomic DNA with primers targeting the TSS or gene body (1kb downstream of TSS) of *JUNB* and *FOSL1*. Each sample was normalized to input control. To calculate a pause index, the occupancy of RNA polymerase II at TSS was divided by the occupancy of RNA polymerase II in the gene body.

### Inhibitors

We used the following inhibitors: Puromycin (Thermo Fisher Scientific), Halt(tm) Protease and Phosphatase Inhibitor Cocktail (Thermo Fisher Scientific), Vemurafenib (Selleck Chemicals), Trametinib (Selleck Chemicals), Dacarbazine (Selleck Chemicals), Temozolomide (Selleck Chemicals), Cisplatin (Selleck Chemicals).

### Antibodies

Cell lysates for immunoblotting were prepared using CelLytic M (Sigma Aldrich) supplemented with a protease and phosphatase inhibitor cocktail, and separated on 4-20% Mini-PROTEAN TGX Precast Protein Gels (Bio-Rad). Proteins were transferred to an Amersham Hybond PVDF membrane (GE Healthcare) using semi-dry transfer, and the binding of HRP-conjugated antibodies was visualized using Clarity Western ECL Substrate (Bio-Rad). For immunoblotting we used the following antibodies: anti-β-actin-HRP clone AC-15 (A3854, Sigma Aldrich), anti-TRIM28 (ab10484, Abcam), anti-FOSL1 (sc-28310, Santa Cruz Biotechnology), anti-JUNB (#3753, Cell Signaling Technologies), anti-YAP1 (#4912, Cell Signaling Technologies), anti-phospho-YAP1 (#4911, Cell Signaling Technologies), anti-ERK1/2 (#9102, Cell Signaling Technologies), anti-phospho-ERK1/2 (#9106, Cell Signaling Technologies), anti-pan-TEAD (#13295, Cell Signaling Technologies) and anti-MITF (#12590, Cell Signaling Technologies). The following secondary antibodies were used: anti-mouse IgG-HRP (#7076, Cell Signaling Technologies), conformation specific anti-rabbit IgG-HRP (#5127, Cell Signaling Technologies), and anti-rabbit IgG-HRP (P0448, Agilent Dako). For ChIP in RNAPII pausing experiments we used anti-RNA polymerase II CTD repeat YSPTSPS antibody (8WG16) (ab817, Abcam) and mouse IgG isotype control (Invitrogen). All antibodies were used at concentrations recommended by the manufacturers.

### Phospho-MAPK array

The phosphorylation of signaling mediators was analyzed using the Human Phospho-MAPK Antibody Array (#ARY002B, R&D Systems) according to the manufacturer’s protocol. Protein lysates were used from A375 and A2058 cells transduced with shSCR or shT28-1 lentivirus. The HRP-coupled streptavidin from the kit was replaced with IRdye 800CW Streptavidin (LI-COR Biosciences), and all signals were analyzed using an Odyssey CLx Imaging System (LI-COR Biosciences).

### Co-immunoprecipitation

Approximately 2×10^7^ A375 cells were lysed in 1 ml of ice-cold lysis buffer (25 mM Tris-HCl pH 7.4, 150 mM NaCl, 1% NP-40, 1 mM EDTA, 5% glycerol) supplemented with a protease and phosphatase inhibitor cocktail. The lysate was rotated slowly at 4°C for 30 minutes. After pre-clearing of the lysate, 1 μg anti-JUNB (#3753, Cell Signaling Technologies) or 1 μg isotype control antibody (#3900, Cell Signaling Technologies) were added to 1.0 mg protein lysate, followed by slow rotation at 4°C for 16 hours. 1.5 mg of magnetic Dynabeads Protein G (Thermo Fisher Scientific) was then added to each lysate, followed by slow rotation at 4°C for 4 hours. Beads were washed three times in ice-cold washing buffer (10 mM Tris-HCl pH 7.4, 150 mM NaCl, 1 mM EDTA, 0.1% Triton X-100) supplemented with a protease and phosphatase inhibitor cocktail. Elution was performed by incubating the beads with SDS-PAGE sample buffer at 50°C for 10 min. Eluates were separated by SDS-PAGE, and detection of co-immunoprecipitated proteins was done by immunoblotting using anti-FOSL1 (sc-28310, Santa Cruz Biotechnology), anti-YAP1 (#4912, Cell Signaling Technologies), or anti-pan-TEAD (#13295, Cell Signaling Technologies), followed by detection with anti-mouse IgG-HRP (#7076, Cell Signaling Technologies) or conformation specific anti-rabbit IgG-HRP (#5127, Cell Signaling Technologies).

### Drug sensitivity assay

Viability assays were performed to measure the response of cells to treatment with Vemurafenib, Trametinib, Temozolomide, Dacarbazine, and Cisplatin. A375 and A2058 cells were seeded in 96-well plates 24 hours prior to the start of the experiment, and then treated for 72-96 hours with drugs. Cell viability was measured using the PrestoBlue Cell Viability Reagent (Thermo Fisher Scientific), and relative viability was calculated as a ratio to untreated control for each condition.

### Gene perturbation with CRISPR/Cas9 and RNAi

TRIM28 knockout cells were generated by transducing A375 cells with LentiCRISPRv2 lentivirus encoding a TRIM28 specific gRNA. After selection with puromycin (1 μg/ml), cells were single-cell sorted into flat bottomed 96-well plates for cell expansion. Loss of TRIM28 expression was verified with anti-TRIM28 immunoblotting. For shRNA mediated knockdown, A375, A2058 and SK-MEL-28 cells were transduced with lentiviruses (LKO.1) that expressed non-targeting control (shSCR or shLuc) or shRNAs specific for TRIM28 (shT28-1 or shT28-2) or YAP1 (shYAP1). After selection in puromycin (1 μg/ml), knockdown efficiency was validated by quantitative RT-PCR and immunoblotting. For CRISPRi, A375 cells were transduced with lentivirus encoding dCas9-KRAB and gRNAs specific for JUNB or EGFP. After selection in puromycin (1 μg/ml), the reduction in expression levels were verified using quantitative RT-PCR and immunoblotting. For siRNA mediated knockdown of JUNB, A375 cells were transfected with siRNA pools targeting JUNB (#L-003269-00-0005, Dharmacon) or non-targeting control siRNA (#D-001810-10-05, Dharmacon). Transfections were performed using Lipofectamine RNAiMAX (Thermo Fisher Scientific) and all cells were transfected with 25 pmol siRNA pools 72 hours prior to analysis.

### Statistical analysis

Statistical analyses were performed using R (R 3.6.0) or Prism 6 (GraphPad Software).

## Data availability

Gene expression data are deposited at the NCBI Gene Expression Omnibus under the accession number GSE133073.

## AUTHOR CONTRIBUTIONS

**WN:** Designed all experiments, performed or contributed to all experiments, analyzed all experiments, performed data analysis, and wrote manuscript. **TH:** Designed, performed and analyzed drug sensitivity experiment. Contributed to analysis of TRIM28 interactome data. **MS:** Contributed to quantitative RT-PCR experiments. Contributed to overexpression experiments. **DVP**: Contributed to Matrigel invasion assays. Contributed to tumor experiments. Contributed to writing the manuscript. **LP**: Generated TRIM28-knockout A375 cells with CRISPR/Cas9. **RC:** Contributed to the analysis of TRIM28 interactome data. Contributed to writing the manuscript. **AE:** Designed all experiments, analyzed all experiments, performed data analysis, and wrote manuscript.

## ACKNOWLEDGEMENTS

This research was supported by the Karolinska Institutet, including KID funding for doctoral education (W.N.) and faculty funded career position (A.E.), Cancerfonden/The Swedish Cancer Society (CAN 2017/680) (A.E.), Vetenskapsrådet/Swedish Research Council (512-2012-1827) (A.E.), Magnus Bergvall foundation (A.E.), Erasmus+ (L.P.). We thank the animal staff at AKM, Center for Molecular Medicine, Karolinska Institutet, for assistance. We acknowledge the staff at the proteomics facility at Karolinska Institutet for assistance with mass spectrometry analysis. The results published here are in part based upon data generated by the TCGA Research Network (https://www.cancer.gov/tcga). The Bioinformatics and Expression Analysis core facility at Karolinska Institutet helped us with performing gene expression analysis. We thank members of Andor Pivarcsi’s laboratory for technical advice. We are grateful to Martin Bergö and Jonas Nilsson for their critical assessment of the manuscript. We thank Marie Wahren-Herlenius for sharing resources and instruments.

## EXTENDED DATA FIGURES

**Extended data Figure. 1.**
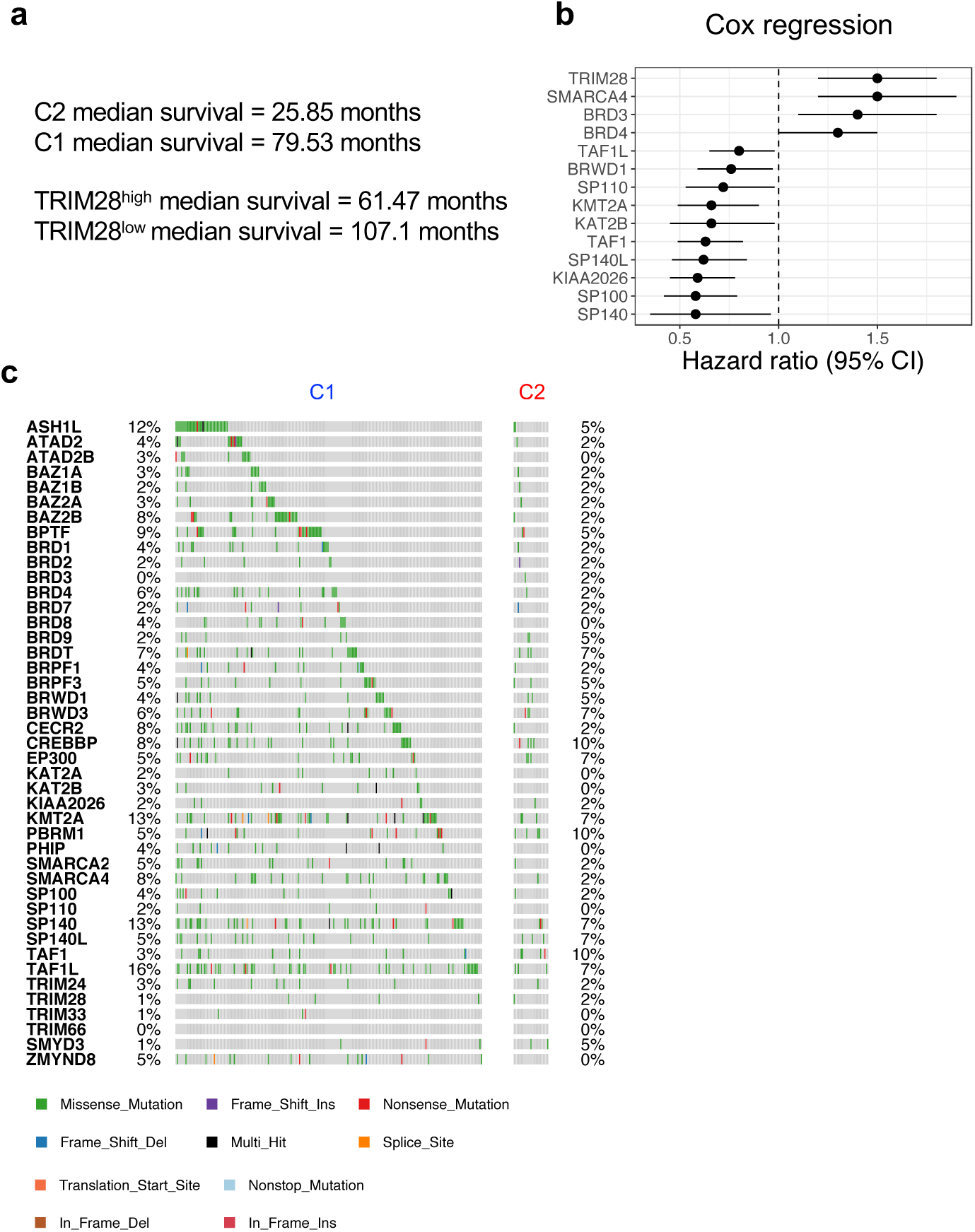
Survival and mutational analysis of patients with cutaneous malignant melanoma. **a**, Median survival of patients in cluster C1 and C2, and of patients with TRIM28^high^ and TRIM28^low^ tumors. RNA-seq from melanoma metastases, and survival data from corresponding patients with cutaneous melanoma, were downloaded from The Cancer Genome Atlas. To obtain stage specific survival we analyzed patients diagnosed with stage III cutaneous melanoma. **b**, Univariate Cox proportional-hazards analysis was performed on bromodomain expression and overall survival in stage III melanoma. To obtain stage specific survival we analyzed patients diagnosed with stage III cutaneous melanoma. RNA-seq from melanoma metastases, and survival data from corresponding patients with cutaneous melanoma, were downloaded from The Cancer Genome Atlas. Univariate Cox proportional-hazards analysis was performed with the *survival* packages in R. Only statistically significant bromodomain genes are displayed. **c**, Whole-exome sequencing data from melanoma metastases was downloaded from The Cancer Genome Atlas. Frequencies of mutations in bromodomain genes from the C1 and C2 tumor clusters were determined using the *maftools* package in R.

**Extended data Figure. 2.**
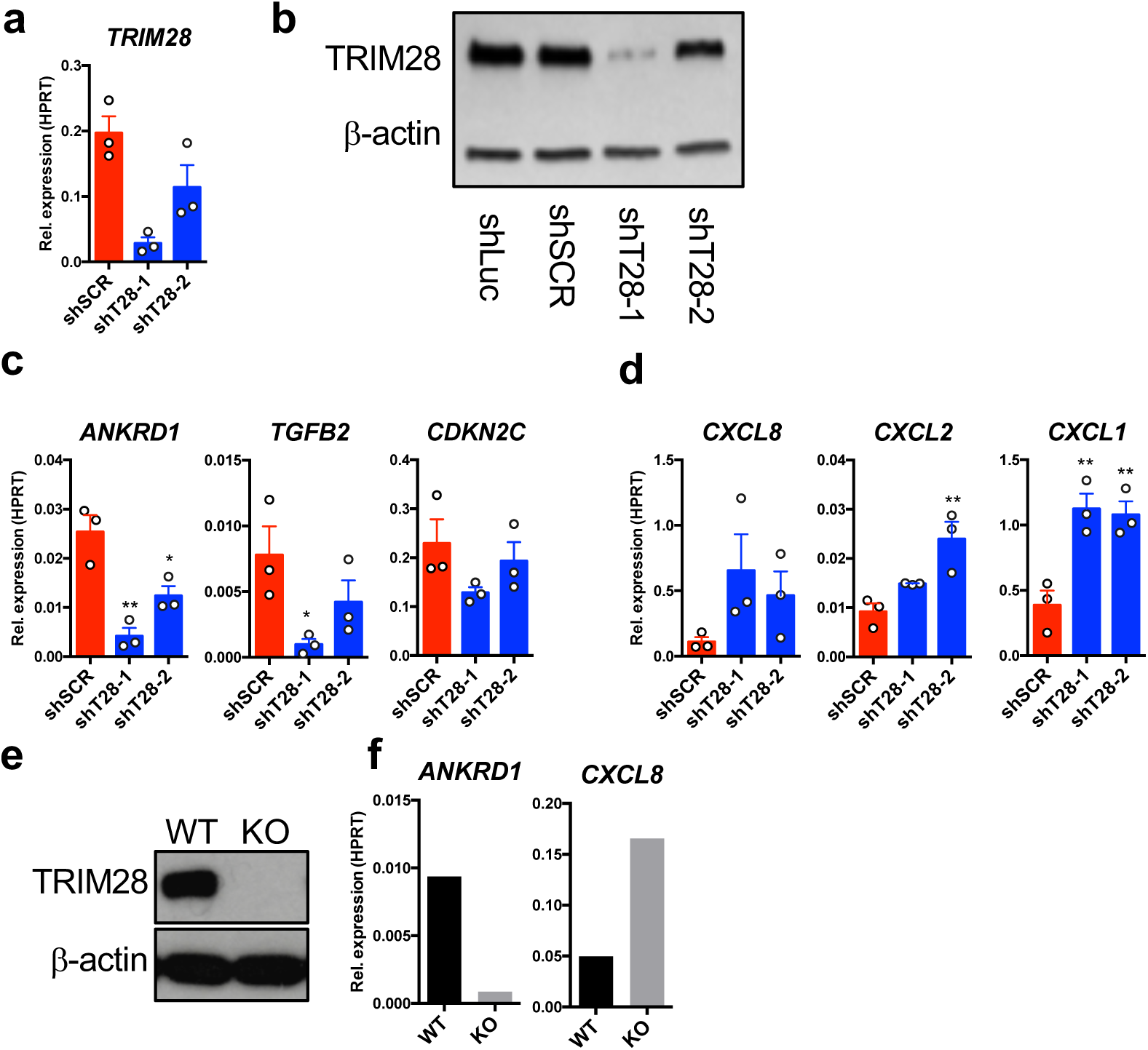
Validation of YAP and KRAS signature genes in A375 cells after knockdown or knockout of TRIM28. **a**, Quantitative RT-PCR validation of TRIM28 knockdown in A375 cells transduced with shSCR, shT28-1 or shT28-2 lentiviruses. Results are expressed as mean ±s.e.m. from three independent experiments (n=3). **b**, Validation of TRIM28 knockdown using immunoblotting on cell lysates from A375 cells transduced with shLuc, shSCR, shT28-1 or shT28-2 lentiviruses. **c**, Quantitative RT-PCR of YAP target genes on A375 cells transduced with shSCR, shT28-1 or shT28-2 lentiviruses. Results are expressed as mean ±s.e.m. from three independent experiments (n=3). One-way ANOVA and Dunnett’s multiple comparison test were used for statistical testing. **d**, Quantitative RT-PCR of *CXCL1, CXCL2* and *CXCL8* on A375 cells transduced with shSCR, shT28-1 or shT28-2 lentiviruses. Results are expressed as mean ±s.e.m. from three independent experiments (n=3). One-way ANOVA and Dunnett’s multiple comparison test were used for statistical testing. **e**, A375 cells were transduced with pLentiCRISPR-v2 containing a TRIM28 specific gRNA. After puromycin selection, single cells were sorted and expanded. Loss of TRIM28 expression was verified using immunoblotting. **f**, Expression levels of *ANKRD1* and *CXCL8* in TRIM28-knockout A375 cells were determined using quantitative RT-PCR. Results are shown as means of technical triplicates.

**Extended data Figure. 3.**
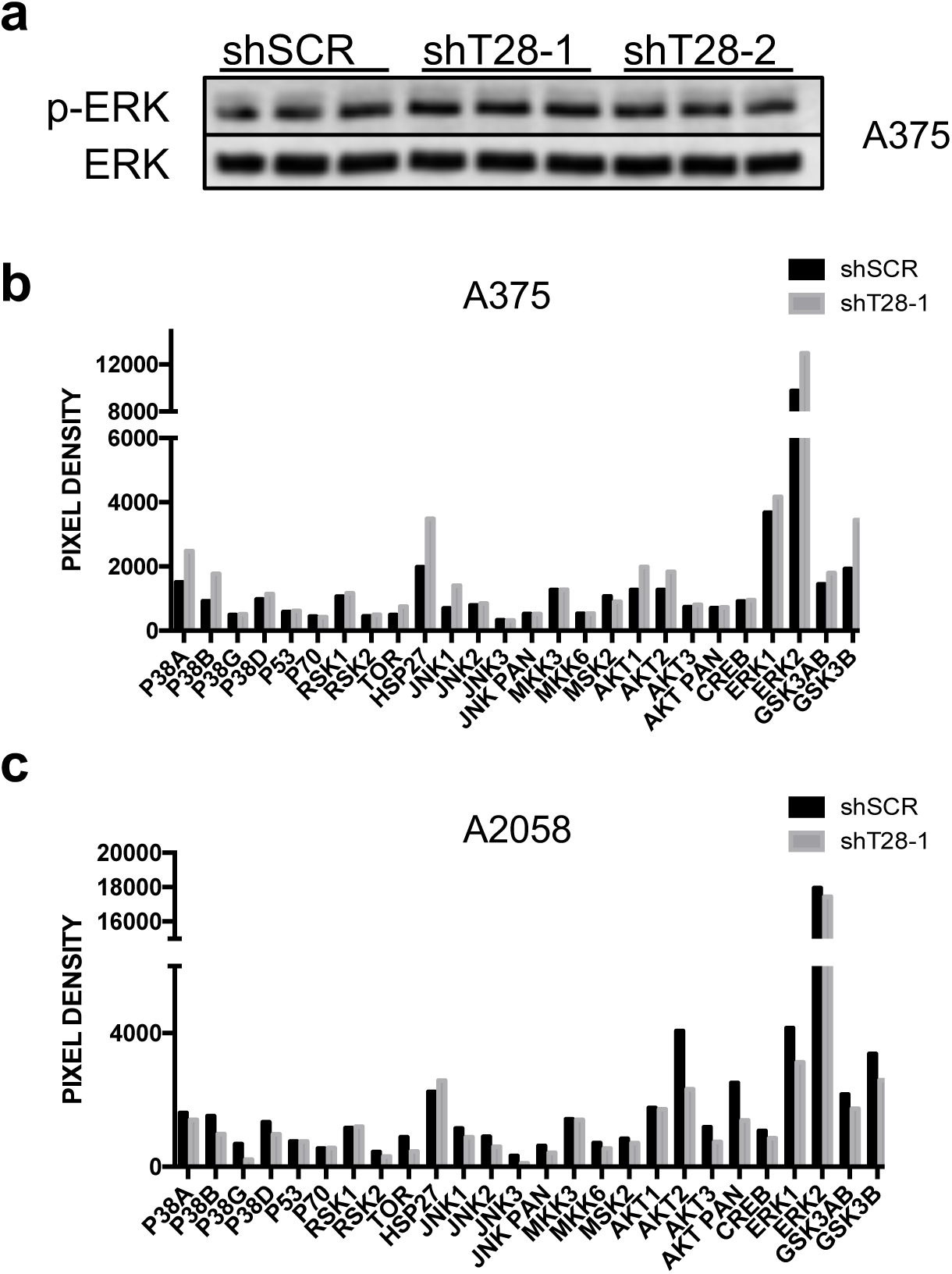
Phospho-profiling of signaling mediators after TRIM28 knockdown in A375 and A2058 melanoma cells. **a**, A375 cells were transduced with shSCR or shT28-1 lentivirus. After selection with 1 μg/ml puromycin cells were lysed and proteins were separated on 4-20% SDS-PAGE. Proteins were transferred to PVDF membranes and proteins were identified with the following antibodies: anti-ERK1/2 and anti-phospho-ERK1/2. **b**, A375 cells were transduced with shSCR or shT28-1 lentivirus followed by selection in medium containing 1 μg/ml puromycin cells. Cells were lysed and lysates were applied to Human Phospho-MAPK Antibody Array. **c**, A2058 cells were transduced with shSCR or shT28-1 lentivirus followed by selection in medium containing 1 μg/ml puromycin cells. Cells were lysed and lysates were applied to Human Phospho-MAPK Antibody Array.

**Extended data Figure. 4.**
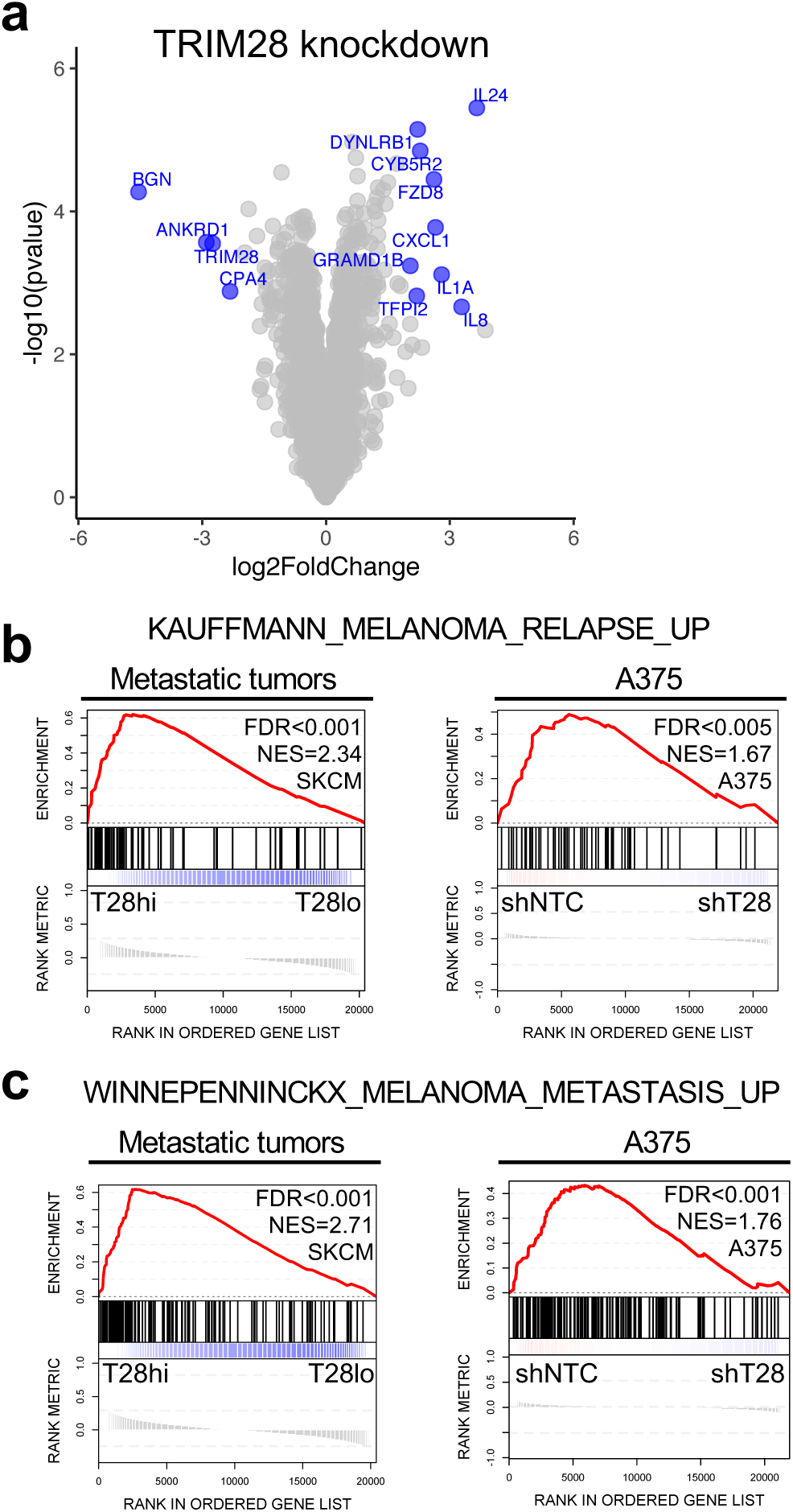
Reduced metastatic gene signatures after TRIM28 knockdown. **a**, A375 cells were transduced with lentiviruses expressing non-targeting control shRNAs (shSCR or shLuc), or TRIM28 specific shRNAs (shT28-1 or shT28-2), followed by selection with puromycin (1 μg/ml). The GeneChip Human Transcriptome Array (HTA) 2.0 (Affymetrix) was used for determination of global gene expression. Significance Analysis of Microarrays (SAM) analysis of gene expression was performed using the R packages *siggenes* and *genefilter.* Volcano plots were generated using the R packages *ggplot2, ggrepel* and *gghighlight*. Genes with a fold change >4 and an adjusted p-value (FDR) <0.05 are highlighted in blue (IL8 = CXCL8). **b, c**, RNA-seq data from melanoma metastases was downloaded from The Cancer Genome Atlas. Gene expression of A375 cells transduced with two TRIM28 specific shRNAs (shT28-1 or shT28-2) and two non-targeting shRNAs (shNTC) was determined using HTA2.0 microarray (Affymetrix). For gene set enrichment analysis (GSEA), expression data files (.gct), phenotype labels (.cls), and gene set files (.gmx) were uploaded to Genomspace (http://www.genomespace.org/) and analyzed using the GSEA tool in GenePattern. False discovery rates were calculated based on gene set permutations. GSEA of a gene signature associated to metastasis of melanoma to distant organs (KAUFFMANN_MELANOMA_RELAPSE_UP) in metastases with high (T28hi) or low (T28lo) TRIM28 expression (left), and in A375 cells transduced with two shNTC or shT28-1 and shT28-2 (right) (**b**). GSEA of a gene signature associated to metastasis of melanoma to distant organs (WINNEPENNINCKX_MELANOMA_METASTASIS_UP) in metastases with high (T28hi) or low (T28lo) TRIM28 expression (left panel), and in A375 cells transduced with two shNTC or shT28-1 and shT28-2 (right panel) (**c**).

**Extended data Figure. 5.**
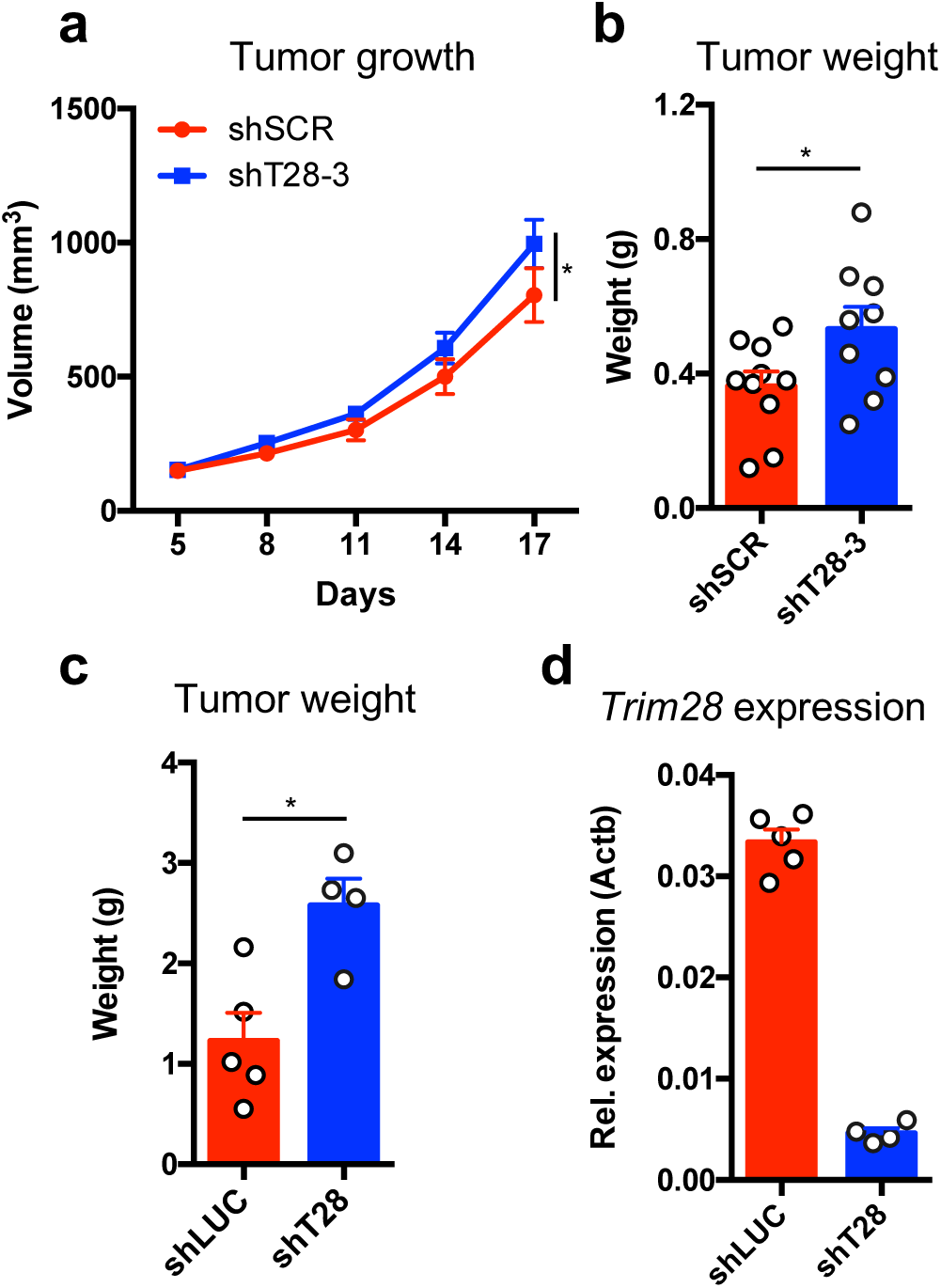
TRIM28 suppresses the growth of melanoma tumors. **a**, Eight-week-old female nude mice were injected subcutaneously with 2.5×10^6^ A375 cells transduced with shSCR or shT28-3 lentivirus in 100 μl of Matrigel diluted to 50% in PBS. Tumor size was measured every third day using a digital caliper and tumor size was calculated using the formula V=(LxWxW)/2. One tumor from shT28-3 transduced A375 cells did not engraft, and was excluded from the analysis. Statistics were calculated using repeated measures ANOVA. **b**, Tumor weight analyzed 17 days after subcutaneous injection of cells transduced with shSCR or shT28-3 lentivirus (n=10 mice per group). Statistics were calculated using the two-sided Mann-Whitney U-test. **c**, Eight-week-old female C57BL6/J were injected subcutaneously with 1×10^5^ B16.F10 cells in 200 μl of Matrigel diluted to 50% in PBS. One tumor from shT28 transduced B16.F10 cells did not engraft, and was excluded from the analysis. After termination of the experiments the tumors were weighed and biopsied for RNA extraction. Tumor weight after subcutaneous injection of 1×10^5^ B16.F10 cells transduced with LKO.1 lentivirus targeting Firefly luciferase (shLuc) (n=5) or Trim28 (shT28) (n=4). Statistics were calculated using the two-sided Mann-Whitney U-test. **d**, Quantitative PCR based validation of *Trim28* gene expression in tumors at the experimental end point.

**Extended data Figure. 6.**
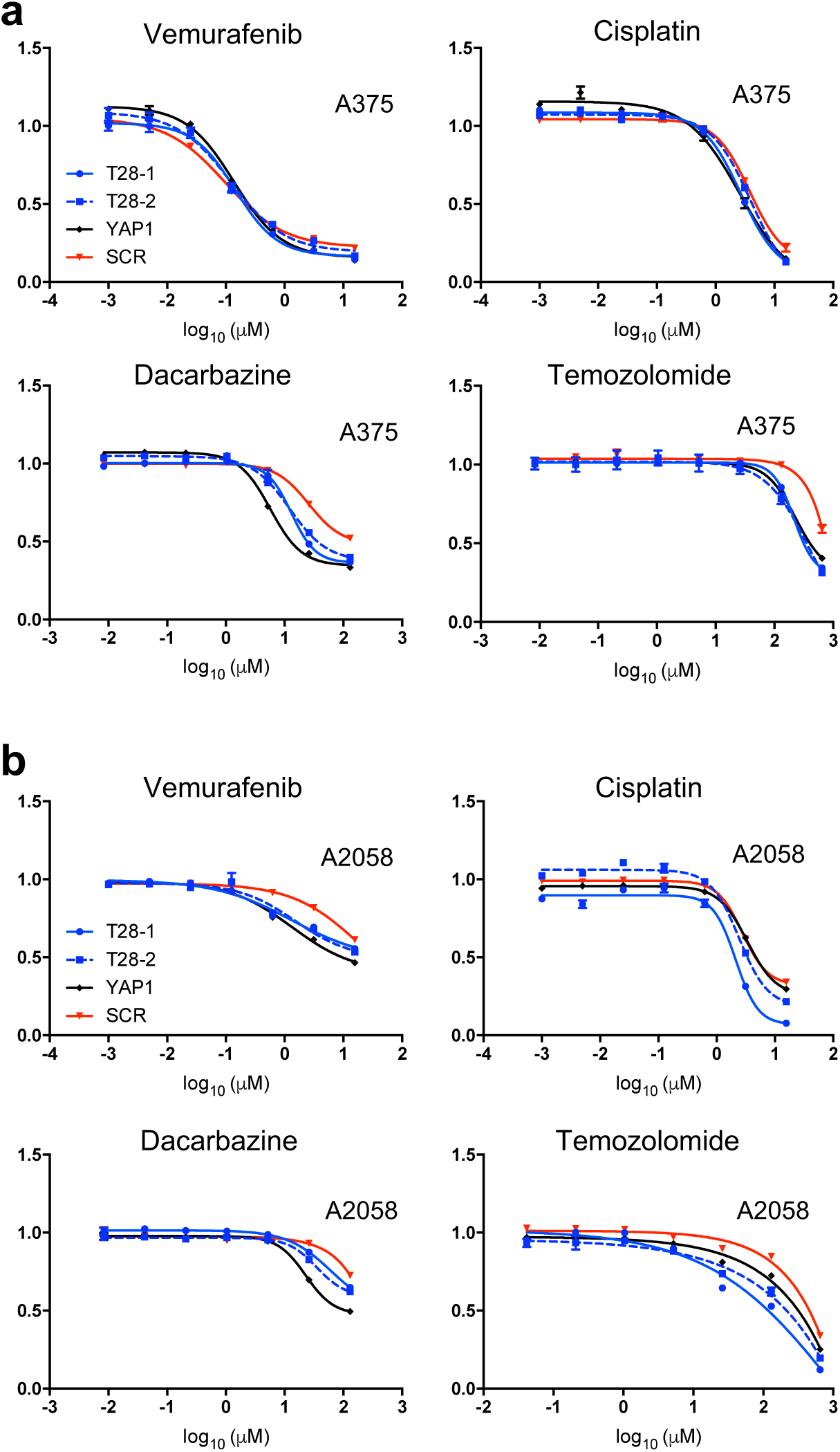
Drug sensitivity of A375 and A2058 melanoma cells after TRIM28 and YAP1 knockdown. **a**, A375 cells transduced with shSCR, T28-1, shT28-2 or shYAP1 lentivirus were seeded in flat-bottomed 96-well plates, and 24 hours later the cells were treated with drug for 72 hours. Viability was determined using PrestoBlue cell viability reagent. One representative experiment of two is shown. **b**, A2058 cells transduced with shSCR, T28-1, shT28-2 or shYAP1 lentivirus were seeded in flat-bottomed 96-well plates, and 24 hours later the cells were treated with drug for 96 hours. Viability was determined using PrestoBlue cell viability reagent. One representative experiment of two is shown.

**Extended data Figure. 7.**
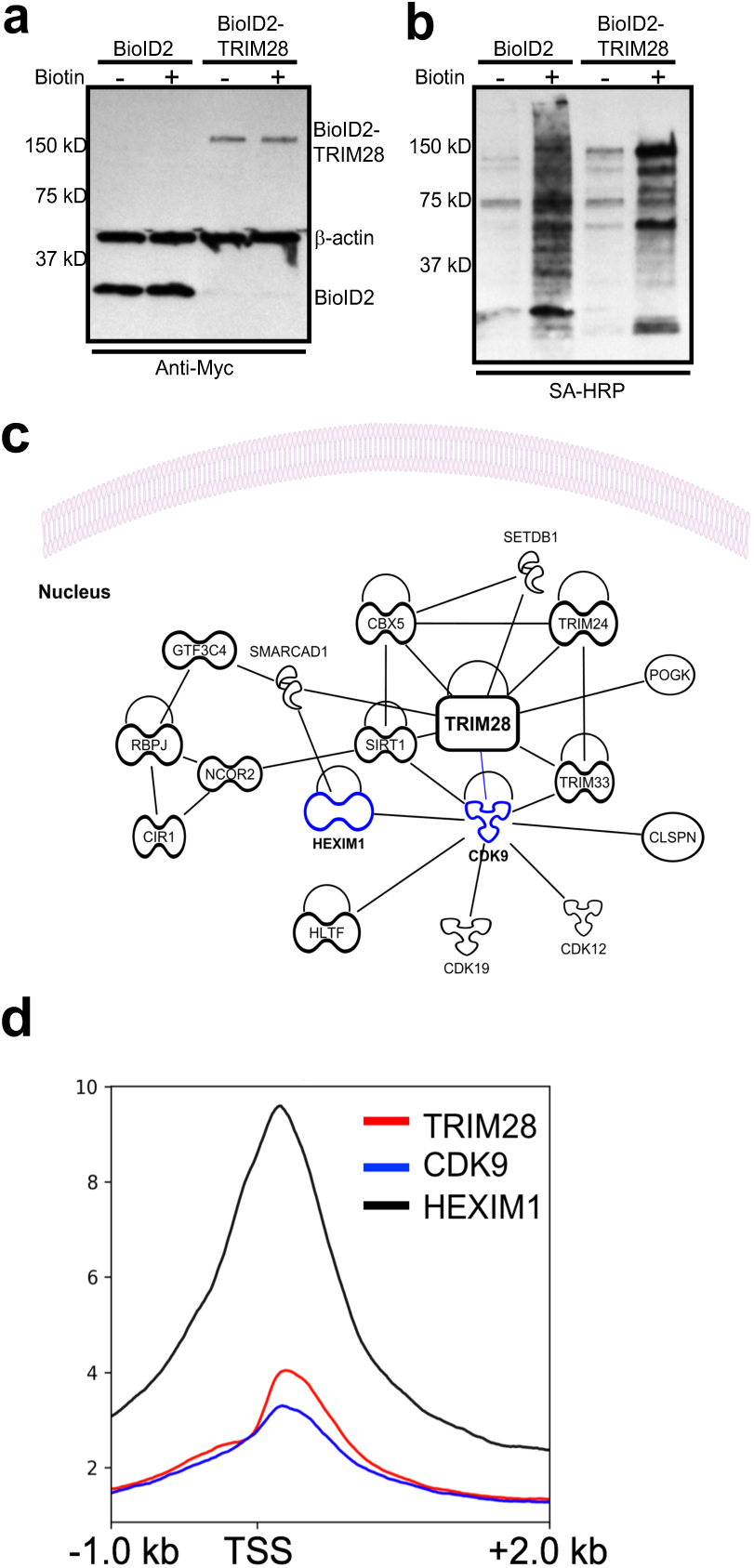
TRIM28 interaction network. **a**, A375 cells were transduced with pBABE-BioID2-TRIM28 or pBABE-BioID2 followed by selection with 1 μg/ml puromycin. Expression of BioID2 and BioID2-TRIM28 in cell lysates was verified with immunoblotting using an anti-Myc antibody. **b**, A375 cells were transduced with pBABE-BioID2-TRIM28 or pBABE-BioID2 followed by selection with 1 μg/ml puromycin. Transduced cells were then cultured in the presence of 50 μM Biotin (Sigma Aldrich) for 20 hours prior to lysis and then bound to Dynabeads MyOne Streptavidin C1 (Thermo Fisher Scientific) magnetic beads overnight. After enrichment of biotinylated proteins, they were detected using streptavidin-HRP. **c**, Protein-protein interaction network based on identified TRIM28 interactors was analyzed by Ingenuity Pathway Analysis (IPA). Displayed interactors are based on experimentally validated direct and indirect interactors. KRAB-ZFN proteins (>50) are not displayed for increased clarity. **d**, ChIP-seq data from HCT-116 cells were analyzed for genome wide co-occupancy of TRIM28, CDK9 and HEXIM1 at transcription start sites (TSS) (GSE72622).

**Extended data Figure. 8.**
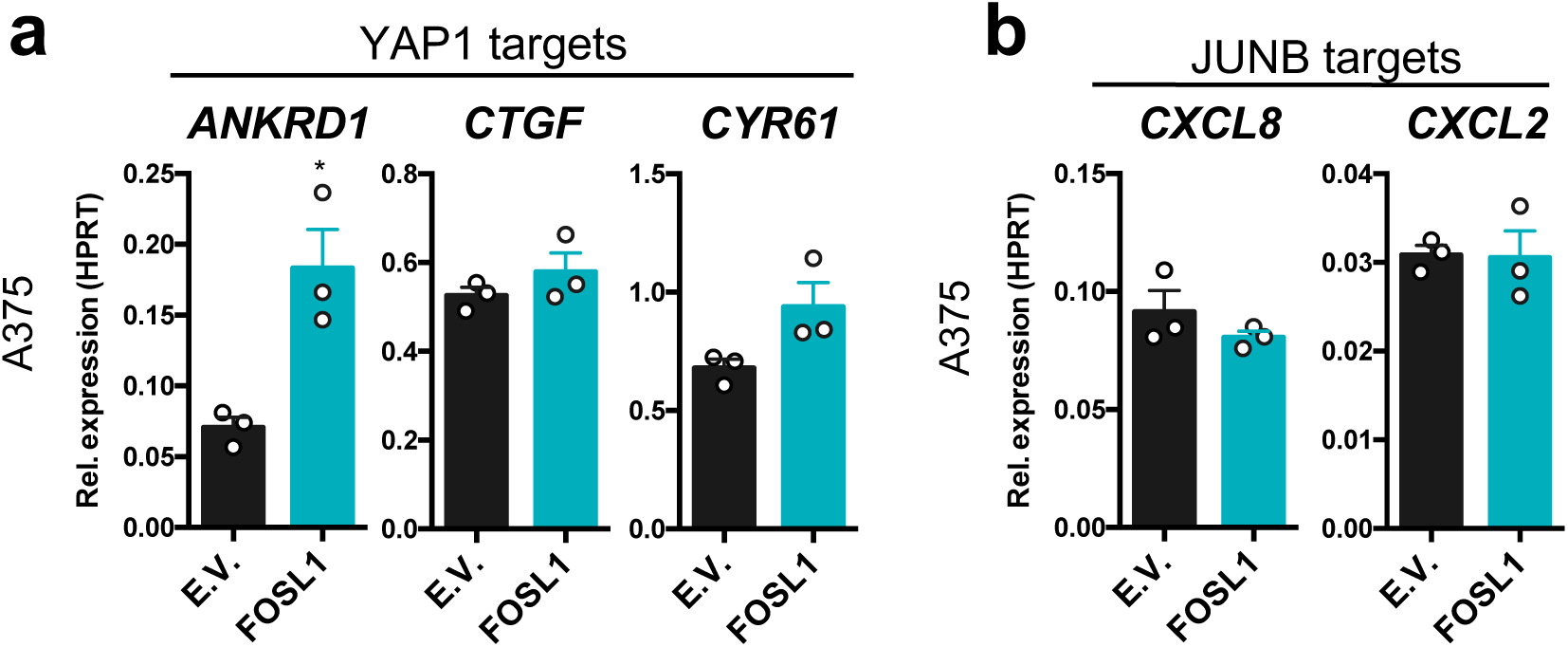
Gene expression after expression of FOSL1. **a**, Expression of YAP1 target genes in A375 cells after transduction with pBABE-FOSL1 or pBABE-E.V. (empty vector). Results are expressed as mean ±s.e.m. from three independent experiments (n=3). Two-sided unpaired t-tests were used for statistical test between the groups. **b**, Expression of *CXCL8* and *CXCL2* in A375 cells after transduction with pBABE-FOSL1 or pBABE-E.V. (empty vector). Results are expressed as mean ±s.e.m. from three independent experiments (n=3). Two-sided unpaired t-tests were used for statistical test between the groups.

**Extended data Figure. 9.**
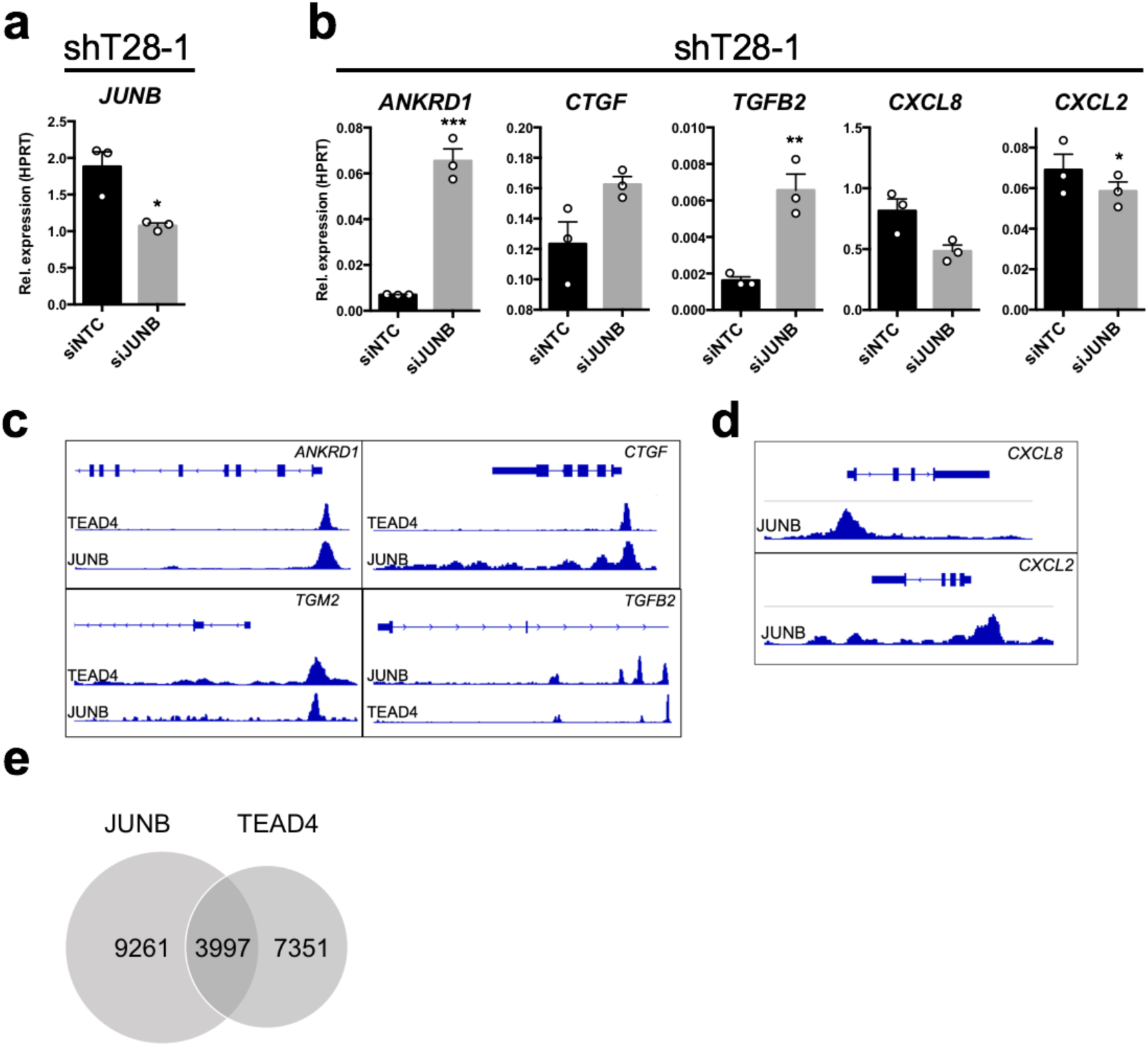
JUNB mediates the effects of TRIM28 knockdown. **a**, Quantitative RT-PCR to determine knockdown efficiency of JUNB in A375 cells previously transduced with shT28-1. Results are expressed as mean ±s.e.m. from three independent experiments (n=3). Two-sided unpaired t-tests were used for statistical test between the groups. **b**, A375 cells, previously transduced with shT28-1, were transfected with siRNA pools of either siJUNB or siNTC. Expression was determined by quantitative RT-PCR. Results are expressed as mean ±s.e.m. from three independent experiments (n=3). Two-sided unpaired t-tests were used for statistical test between the groups. **c**, ChIP-seq analysis of co-occupancy of JUNB and TEAD4 at promoters of canonical YAP1 signature genes in A549 cells. **d**, ChIP-seq analysis of JUNB occupancy at promoters of the canonical KRAS signature genes *CXCL2* and *CXCL8* in A549 cells. **e**, Genome-wide overlap of ChIP-seq peaks for JUNB and TEAD4 in A549 cells. ChIP-seq data in **(c-e)** are from GSE32465 and GSE92807.

**Extended data Figure. 10.**
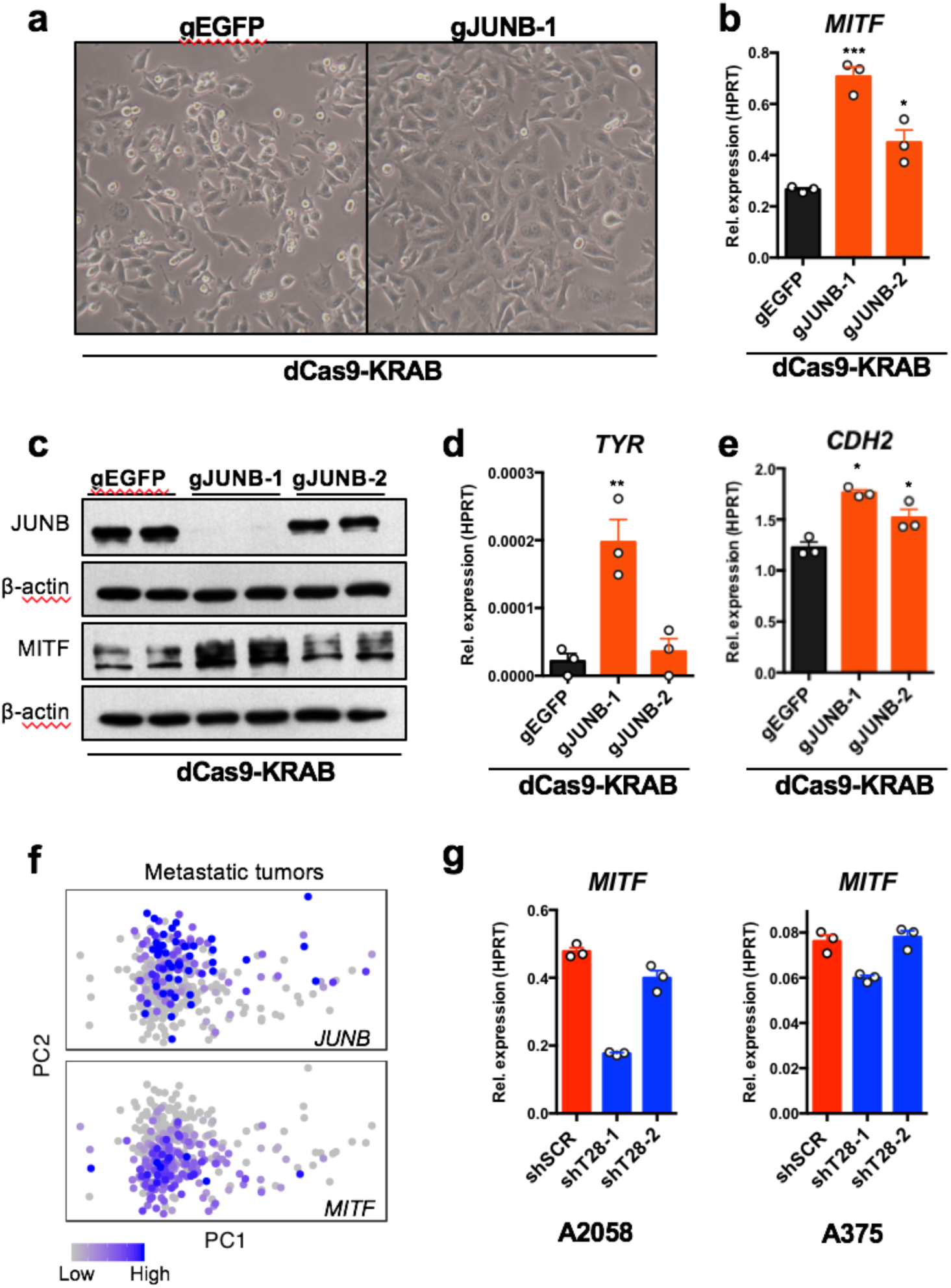
JUNB controls the expression of MITF and N-cadherin in melanoma cells. **a**, Images displaying the morphology of A375 cells transduced with dCas9-KRAB lentiviruses expressing a control gRNA (gEGFP) or a gRNA targeting *JUNB* (gJUNB-1). **b**, Expression of *MITF* after transduction of A375 cells with dCas9-KRAB lentiviruses expressing gRNA targeting *JUNB* (gJUNB-1 or gJUNB-2) or a control gRNA (gEGFP). Expression was determined by quantitative RT-PCR. Results are expressed as mean ± s.e.m. from three independent experiments (n=3). One-way ANOVA and Dunnett’s multiple comparison test were used for statistical testing of differences to gEGFP. **c**, Immunoblotting against JUNB and MITF on protein lysates from A375 cells after transduction with dCas9-KRAB lentiviruses expressing gRNA targeting *JUNB* (gJUNB-1 or gJUNB-2) or a control gRNA (gEGFP). **d**, Expression of the MITF target gene *TYR* after CRISPRi against JUNB. Expression was determined by quantitative RT-PCR. Results are expressed as mean ± s.e.m. from three independent experiments (n=3). One-way ANOVA and Dunnett’s multiple comparison test were used for statistical testing of differences compared to gEGFP. **e**, Expression of N-cadherin (*CDH2*) after transduction of A375 cells with dCas9-KRAB expressing gRNAs targeting *JUNB* (gJUNB-1 and gJUNB-2) or a control gRNA (gEGFP). Expression was determined by quantitative RT-PCR. Results are expressed as mean ±s.e.m. from three independent experiments (n=3). One-way ANOVA and Dunnett’s multiple comparison test were used for statistical testing of differences compared to gEGFP. **f**, Expression levels of *JUNB* and *MITF* indicated for patient derived metastatic tumors (n=367) and displayed in a principal component analysis plot. Results are based on RNA-seq data from TCGA. **g**, Expression of *MITF* after transduction of A375 cells with shSCR, shT28-1 or shT28-2. Expression was determined by quantitative RT-PCR.

